# Task-specific theta enhancement and domain-general alpha/beta suppression as oscillatory signatures of individual differences in cognitive flexibility

**DOI:** 10.1101/2025.09.26.678741

**Authors:** Mareike J. Hülsemann, Christoph Löffler, Anna-Lena Schubert

## Abstract

Cognitive flexibility—the ability to adapt to changing situational or task demands—is a fundamental aspect of human cognition and a potential source of individual differences in higher-order cognitive abilities. To further understand the neural mechanisms underlying this ability, we examined event-related spectral perturbations (ERSP) in three cued switching tasks (parity/magnitude, global/local, number/letter) in a sample of 148 adults. Employing a data-driven approach that combined mass-univariate time-frequency analysis, cluster-based permutation testing, and latent change structural equation modeling, we identified two robust neural signatures: a transient frontal theta increase and a sustained alpha/beta suppression. Both effects were more pronounced in switch compared to repeat trials. Frontal theta reflected task-specific control processes, whereas parietal alpha/beta indexed domain-general attentional processes that persisted from cue onset through target processing—consistent with two stage-models of task switching. Notably, flexibility-related alpha/beta dynamics did not correlate with measures of intelligence or working memory capacity (WMC), underscoring the distinctiveness of cognitive flexibility from general cognitive abilities. These findings provide the first direct evidence that alpha/beta suppression constitutes a reliable, generalizable neural signature of individual differences in cognitive flexibility.

## Introduction

Cognitive flexibility is a key element of human behavior that enables us to adapt to changing circumstances—whether switching between different tasks during the working day (Koch, Poljac, Müller, & Kiesel, 2018) or generating creative solutions to novel problems (Palmiero, Fusi, Crepaldi, Borsa, & Rusconi, 2022). We define cognitive flexibility as the ability to adapt to changing task demands by switching between mental sets or rules (Hohl & Dolcos, 2024). As a core component of executive function models, alongside in-hibition and updating, cognitive flexibility is theorized to underlie more complex goal-directed processes such as planning and problem-solving (Miyake et al., 2000). Yet while cognitive flexibility represents a fundamental capacity for adaptive behavior, individuals show marked differences in their flexibility abilities, and understanding these individual differences remains a central challenge in cognitive psychology.

### Individual differences in cognitive flexibility

Individual differences in cognitive flexibility have profound implications across multiple domains of human functioning. They predict meaningful real-world outcomes such as academic achievement (Cortés Pascual, Moyano Muñoz, & Quílez Robres, 2019; Zheng, Akaliyski, Ma, & Xu, 2024), problem-solving capabilities (Bobrowicz & Thibaut, 2023), social interactions (Harel, Hemi, & Levy-Gigi, 2023), and mental health outcomes (Dajani & Uddin, 2015). Conversely, reduced cognitive flexibility characterizes numerous clinical populations, including individuals with schizophrenia (Laere, Tee, & Tang, 2018), depression (Dotson et al., 2020), and autism spectrum disorder (Uddin, 2021). Some researchers propose that individual differences in cognitive flexibility may even explain the positive correlations observed across diverse cognitive tasks, as most complex cognitive operations require some degree of flexible adaptation (Kovacs & Conway, 2016; Santarnecchi et al., 2021).

### Neural mechanisms underlying cognitive flexibility

To understand individual differences in cognitive flexibility, researchers have turned to neural measures that can provide mechanistic insights beyond behavioral performance alone. Cognitive flexibility is typically assessed using a cued task-switching paradigm, where participants must switch between different rulesets based on contextual cues (Miyake et al., 2000; Gratton, Cooper, Fabiani, Carter, & Karayanidis, 2018). This paradigm reveals cognitive flexibility through comparing switch trials (requiring rule changes) to repeat trials (maintaining the same rule), with flexibility indexed by switch costs in reaction time and accuracy.

Electrophysiological studies using event-related potentials (ERPs) and time-frequency analyses have identified consistent neural signatures of cognitive flexibility. ERP studies show greater centro-parietal positivity during cue processing in switch versus repeat trials, reflecting proactive control processes, and reduced P3b amplitude following target presentation in switch trials, suggesting ongoing reconfiguration demands (Karayanidis & Jamadar, 2014). In addition, oscillatory analyses reveal robust spectral signatures: enhanced theta power increases and alpha power suppression are consistently observed in switch compared to repeat conditions (Cunillera et al., 2012; Cooper, Wong, McKewen, Michie, & Karayanidis, 2017; Barlow, Medrano, Seichepine, & Ross, 2018; Capizzi, Ambrosini, Arbula, & Vallesi, 2020). Frontal theta enhancement reflects active cognitive control and working memory recruitment (Cavanagh & Frank, 2014; Sauseng, Griesmayr, Freunberger, & Klimesch, 2010), while parietal alpha suppression indicates disengagement from task-irrelevant information (Van Diepen, Foxe, & Mazaheri, 2019; Sauseng et al., 2006). Crucially, these neural signatures show stability over time (Cooper et al., 2019) and scale with task difficulty (Wu et al., 2015), suggesting they reflect meaningful individual differences in cognitive control capacity.

### Critical knowledge gap: Neural correlates of individual differences

Despite this rich understanding of both behavioral individual differences and neural mechanisms at the group level, a fundamental disconnect remains: we do not yet understand whether individual variation in neural signatures like theta enhancement and alpha suppression accounts for the behavioral variability observed across participants. This represents a critical gap in our mechanistic understanding of cognitive flexibility. Moving beyond group-average neural responses to examine individual differences in these neural markers is essential for developing a precise understanding of how brain oscillations support flexible cognition (Braver, Cole, & Yarkoni, 2010).

However, investigating neural correlates of individual differences requires establishing that neural measures meet basic psychometric standards. Currently, the reliability of neural flexibility measures has received insufficient attention in cognitive neuroscience (Clayson, 2024), despite reliability being a prerequisite for valid individual differences research (Gell et al., 2024). Without reliable neural measures, variations in brain signals cannot be confidently attributed to genuine individual differences rather than measurement noise. Furthermore, for neural measures to serve as meaningful individual difference variables, they must demonstrate convergent validity across different flexibility tasks and criterion validity through associations with relevant psychological constructs such as working memory capacity (WMC) and fluid intelligence.

Addressing this knowledge gap requires overcoming several methodological limitations that have constrained previous research on neural mechanisms of cognitive flexibility. First, some studies use simultaneous cue-target presentation or very short cue-target intervals, which conflates endogenous preparation processes with exogenous implementation processes (Rogers & Monsell, 1995; Kiesel et al., 2010). Understanding individual differences requires separating these temporally distinct control processes, as individuals may vary in their capacity for endogenous versus exogenous control strategies.

Second, many previous studies have focused on predefined frequency bands and brain regions, typically examining theta and alpha activity over frontal and parietal regions (Cunillera et al., 2012; Foxe, Murphy, & De Sanctis, 2014). While this approach has identified important effects, it may miss other frequency bands or brain regions that contribute to individual differences in flexibility. A data-driven mass-univariate approach with cluster-based permutation testing (Groppe, Urbach, & Kutas, 2011) offers greater statistical power (Fields & Kuperberg, 2020) and enables assessment of effect specificity across the full time-frequency-space domain.

Third, most research examines single tasks, despite evidence that substantial variance in flexibility measures reflects task-specific rather than general flexibility processes (Löffler, Frischkorn, Hagemann, Sadus, & Schubert, 2024). Individual differences in general cognitive flexibility can only be validly assessed by extracting common variance across multiple tasks, mathematically eliminating task-specific influences such as processing speed, semantic proficiency, or stimulus-specific effects.

Finally, previous studies often used modest sample sizes that preclude robust individual differences analyses and limit the reliability of neural measures. Adequate statistical power is essential for detecting individual differences in neural oscillations and for implementing advanced analytical approaches such as structural equation modeling.

### The present study

The present study addresses these methodological challenges to provide the first comprehensive investigation of individual differences in neural correlates of cognitive flexibility. We employed three distinct switching tasks with relatively long cue-target intervals (800–1200ms) to distinguish endogenous and exogenous control processes. Using a data-driven timefrequency approach across the full scalp, we identified neural signatures of flexibility and assessed their reliability and validity as individual difference measures. Most importantly, by examining common variance across multiple flexibility tasks in a large sample (N = 148), we isolated neural correlates of general cognitive flexibility while eliminating task-specific influences.

This methodologically rigorous approach enables us to answer fundamental questions about the neural basis of individual differences in cognitive flexibility: Are these neural individual differences reliable across time and valid across tasks? And do neural flexibility measures relate to other cognitive abilities in theoretically meaningful ways? By bridging the gap between neural mechanisms and individual differences research, this study has the potential to advance both theoretical models of executive functioning and applied research on assessment of cognitive control skills.

## Methods

### Sample

The final sample consisted of 148 participants from the general population (96 females, 51 males, one person declared no affiliation to either gender). The median age of participants was 24 years, with an age range of 18 to 60 years (*M* = 31.5, *SD* = 13.9). All participants were fluent in German, with all but four being native speakers. Thirteen participants stated to be left-handed. A total of 38.5 % of the sample had obtained a university degree, while 58.8 % had received a high school diploma as their highest qualification. The majority of the sample (58.8 %) was currently enrolled in a university program. The sample was recruited via advertisements in local newspapers, flyers, and via the departmental participant pool of Heidelberg university. Initially, 151 people were invited to participate in the study. However, three participants subsequently withdrew from the study. Participants who suffered from claustrophobia, were undergoing psychiatric or neurological treatment, had psychological or neurological disorders, were taking medication that affected the nervous system, had skin allergies, had a pacemaker, were colour-blind, were obese, or had chronic back pain were excluded. Participants provided written informed consent prior to participation. For participation, they received monetary compensation and personalized feedback on their performance on intelligence and working memory tests. The study was approved by the local ethics committee (reference number: Löf 2019/1–3) and is in accordance with the Declaration of Helsinki (World Medical Association, 2013).

### Openness and transparency

This data set is available via OSF (https://osf.io/4pvz3/) and was initially described in (Löffler, Frischkorn, et al., 2024). This comprehensive dataset has been analyzed and reported on in previous publications (Schubert, Löffler, Jungeblut, & Hülsemann, 2025; Schubert, Frischkorn, et al., 2024; Schubert, Löffler, Wiebel, Kaulhausen, & Baudson, 2024; Sadus, Schubert, Löffler, & Hagemann, 2024; Löffler, Sadus, Frischkorn, Hagemann, & Schubert, 2024; Lesche, Sadus, Schubert, Löffler, & Hagemann, 2024; Schubert, Löffler, & Hagemann, 2022; Schubert, Löffler, Hagemann, & Sadus, 2023). Access to the preprocessed data and analysis code used in this paper is provided via Github (https://github.com/mhuelsemann/flexibility_TFA). The study and current analyses were not pre-registered.

### Materials

All computer-based tasks were programmed in MATLAB 2018b (The MathWorks Inc., Natick, Massachusetts) using Psychtoolbox version 3.0.13 (Kleiner, Brainard, & Pelli, 2007). They were presented on a 21.3” LCD monitor (Eizo FlexScan S2100) with a 1280 × 1024 resolution and a 60 Hz refresh rate. Participants had a distance of approximately 100 cm to the computer screen.

### Measurement of Flexibility

We used the well-validated and widely used task-cueing paradigm to investigate cognitive flexibility (Kiesel et al., 2010). We implemented three different shifting tasks, which require participants to switch between two categorization tasks indicated by a cue. In each trial, participants had to classify the target according to the cue. The color of the cue (fixation cross in red or green) indicated which rule-set to use. The cue was always valid, such that cue and target had the same color throughout all trials.

For all tasks, a trial consisted of a sequence of four slides (Figure 1a). First the cue was presented 400 to 600 ms (randomly selected from a uniform distribution). The cue was followed by a blank screen for 400 to 600 ms (randomly selected from a uniform distribution), referred to as inter stimulus interval (ISI). Then the target was presented until the participant responded, but always for a minimum of 1000 ms and a maximum of 3000 ms. Lastly, there was an inter trial interval (ITI) of 1000 to 1500 ms (randomly selected from a uniform distribution), showing again a blank screen. The duration of one trial varied between 2800 and 5700 ms, depending on cue presentation time, ISI and ITI intervals, as well as participants’ response time (i.e. target presentation time).

**Figure 1.**
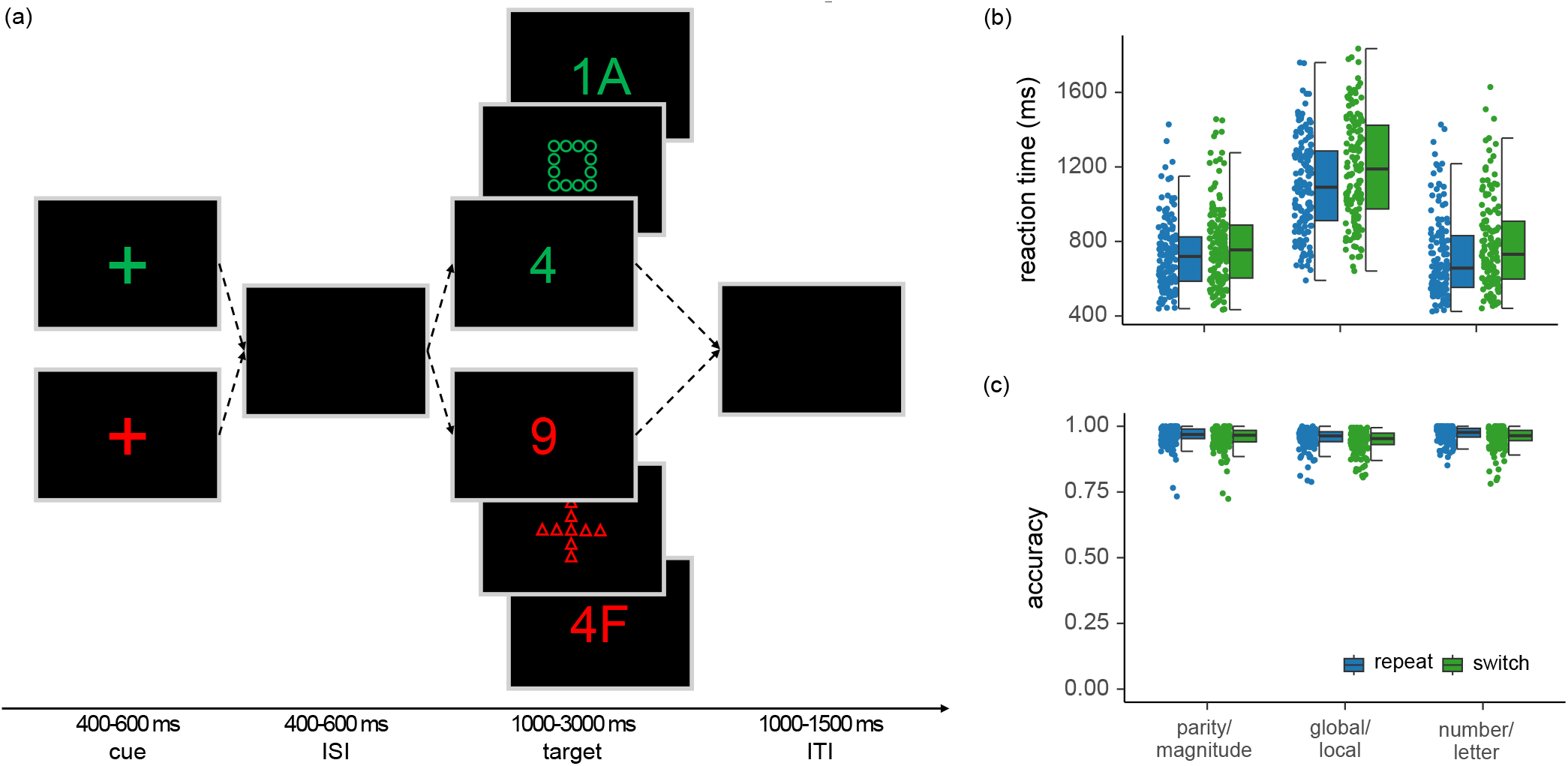
A schematic illustration of a trial of the cued task-switching paradigm and behavioral results. *Note*. (a) Each trial starts with a colored fixation cross, cueing the subsequent rule-set. After a variable inter-stimulus interval, the target was presented in the same color as the cue. Participants had to respond according to the correct rule-set by a key press. Exemplary targets for each of the three shifting tasks are illustrated (from inner to outer: parity/magnitude, global/local, number/letter). (b) Mean reaction times (ms) and (c) accuracy (%) for all three shifting tasks for both conditions are depicted.

In the *parity/magnitude task*, participants had to decide whether the target, a number between one and nine (except five), was either smaller or larger than five (ruleset magnitude, red) or whether it was odd or even (ruleset parity, green). This task was adapted from Sudevan and Taylor (1987) and had total of 384 trials.

In the *global/local task*, the target was one of four geometric figures (circle, triangle, square, cross). Each figure was composed of small geometric shapes from the same set. The larger figure and the smaller shapes could never have the same geometrical shape. Participants had to indicate what shape a figure was (rule-set global, red) or what shapes it was made of (rule-set local, green). This task was adapted from Miyake et al. (2000) and had total of 384 trials.

In the *number/letter task*, the target was a number-letter pair (e.g., 3K). Participants had to decide whether the presented number was smaller or larger than five (rule-set number, red) or whether the letter was a vowel or consonant (rule-set letter, green). Numbers from one to nine (except five) and letters A, E, I, U, G, K, M, and R were used. This task was adapted from Rogers and Monsell (1995) and a total of 256 trials were presented.

Of all trials in each switching task, 50 % were switch trials, requiring a shift in the rule-set, and 50 % were repeat trials, requiring the same rule-set as in the trial before. Half of all trials belonged to each rule-set. Condition (switch vs. repeat) and rule-set were fully crossed. Trial conditions were randomized with the restriction that neither rule-set nor condition occurred more than three times in a row. Furthermore, none of the targets occurred twice in a row. Mapping of response keys was kept constant across participants.

Before starting the experimental trials, participants practiced the task, receiving feedback on the correctness of their response. The practice consisted of 10 trials for each rule-set followed by 20 trials, in which both rule-sets were intermixed. Participants were instructed to respond as fast and accurate as possible, by pressing one of two keys (parity/magnitude task and number/letter task) or one of four keys (global/local task) on the keyboard. If participants did not respond within 3000 ms of target onset, the trial was marked as missing and during practice participants received the feedback “too slow”.

Slides were black with either red (#FF1920) or green (#00B21E) stimuli, depending on the rule-set. Numbers and letters were presented in typeface Arial and font size 40. All stimuli were centered horizontally and vertically.

As expected, and consistent with previous research, we observed longer reaction times in switch trials than in repeat trials across all three switching tasks. See Figures 1b and 1c for reaction times and accuracy, respectively.

### Intelligence

To assess fluid intelligence, we used the short version of the Berlin Intelligence Structure Test (BIS, Jäger, Süß, & Beauducel, 1997). Here, a total of 15 tasks were utilized to fully cross the four operation-related components of intelligence (processing capacity [PC], processing speed [PS], memory [M], and creativity [C]) with the three content-related components of intelligence (verbal, numerical, and figural). The short version of the BIS was completed in approximately 50–60 minutes and was conducted in groups of up to 4 participants. The operation-related component scores for each participant were calculated by aggregating the normalized *z*-scores of all tasks that measured the respective component. Participants had a mean IQ of 96 (*SD* = 16).

### Working memory capacity

To evaluate the WMC of the participants, the Working Memory Test Battery, developed by (Lewandowsky, Oberauer, Yang, & Ecker, 2010), was employed. Four tasks from the battery were utilized: the memory updating task, the operation span task, the sentence span task, and the spatial short-term memory task. However, due to an error in the programming code, we could not use the data of the spatial short-term memory task in our analyses. Furthermore, we employed the location-letter binding task, as described by (Wilhelm, Hildebrandt, & Oberauer, 2013). With the exception of five participants, all subjects completed this letter binding task. For each set size in the working memory tasks, the mean proportion of correctly solved items was calculated for each participant as the dependent variable.

### Procedure

The study comprised three measurement occasions, with data collected at three-month intervals. The duration of each measurement occasion was approximately 3.5 hours. Prior to their inclusion in the study, participants were required to sign an informed consent form at the beginning of the first session. They then completed the Ishihara test (Ishihara, 1972) to ascertain that they exhibited no form of color blindness, as this could have affected the results of the study. In the initial two sessions, an electroencephalogram (EEG) was recorded while participants completed six distinct cognitive tasks on a computer (for details see Löffler, Frischkorn, et al., 2024). EEG was recorded in a dimly lit, electromagnetically shielded cabin. To minimize between-subjects error variance, the tasks were presented in a consistent sequence for the entire sample (Goodhew & Edwards, 2019). In the first session, participants completed the parity/magnitude task and the global/local task. Additionally, demographic data about the participants was gathered at the end of the first session. In the second session, the number/letter task was administered. The third session was conducted in groups of 1 to 4 participants. First, the BIS was administered as a paper-and-pencil test. Subsequently, the participants completed the WMC tasks on computers. Furthermore, participants completed two brief cognitive tests measuring cognitive abilities, a questionnaire assessing mindwandering, and a pretzel task (the results of which are not presented here).

### EEG recording and quantification

#### Hardware

EEG was continuously recorded with 32 equidistant passive silver-silver chloride (Ag/AgCl) electrodes (EasyCap GmbH, Herrsching, Germany). See Figure A1 in the appendix for the electrode layout. All sites were referenced online to Cz. Fpz served as ground. We used a BrainAmp DC amplifier with an input impedance of 10*M*Ω (Brain Products GmbH, Munich, Germany). Recordings, in AC mode, were sampled at 1,000 Hz with a 0.1*µV* resolution. The impedances of EEG electrodes were kept below 5*k*Ω. The pass-band of the hardware filter was set to 0.016 to 1000 Hz. Due to recordings errors, data from two subjects in session one and four subjects in session two had to be discarded. EEG data was processed in MATLAB (R2021a, The MathWorks, Inc.) using the EEGLAB 2023.0 toolbox (Delorme & Makeig, 2004).

#### Pre-processing for independent component analysis

EEG data was analyzed independently for each task. We segmented the data from the beginning to the end of each task and then performed the following preprocessing and data analysis steps. Raw EEG data was first pre-processed for running independent component analysis (ICA, Bell & Sejnowski, 1995). ICA was used for removing eye, muscle, and heart artefacts. Data was high pass filtered at 1 Hz using a zerophase Hamming windowed sinc FIR filter implemented in EEGLAB (Widmann, Schröger, & Maess, 2015). Noisy channels were rejected using the channel criterion of the clean_artefacts function (Kothe, Miyakoshi, & Delorme, n.d.). Channels were rejected if they correlated less then *r* = .80 to their neighboring channels. Across all tasks, on average 0.60 ± 0.83 (median = 0) channels were removed in each data set ranging from 0 to 4. Line noise was removed using Cleanline (v2.0) with the default settings (Mullen, 2012; Bigdely-Shamlo, Mullen, Kothe, Su, & Robbins, 2015). Then the data was re-referenced to the average reference and the online reference Cz was recovered. Cz was then discarded after re-referencing to match the number of channels to the rank of the data for ICA. Cz was chosen because its activity is best captured by neighboring electrodes. Therefore, presumably the least amount of information is lost by discarding this electrode. Data was temporally segmented into consecutive two-second epochs and artefactual epochs were rejected automatically using threshold, joint probability, and kurtosis (Delorme, Sejnowski, & Makeig, 2007). Epochs were rejected if activation exceeded amplitude thresholds of ± 1000*µV*. Furthermore, improbable data epochs were discarded on the assumption that epochs with artifacts are improbable. This was done by computing the probability distribution of all data points, deriving the probability of each data point, and defining the probability of an epoch by the joint probability of all data points within an epoch. Probability was calculated for single electrodes and for the entire set of electrodes. Improbability was determined by the standard deviation of the mean probability distribution. The threshold was set at five *SD* for single channels and two *SD* for all channels. Similarly, epochs were classified as artefactual if the kurtosis of the data point distribution within an epoch exceeded a threshold. Again, the threshold, determined as the standard deviation from the mean kurtosis values, was set at five *SD* for single channels and two *SD* for all channels. Rejection of an epoch always led to the rejection of this epoch in all channels. This automatic procedure led to an average of 12 ± 3 % of rejected epochs (range: 5 – 20 %) across all tasks. ICA was performed on the remaining segments, using the extended infomax algorithm (Bell & Sejnowski, 1995; Lee, Girolami, & Sejnowski, 1999). ICs were automatically classified using ICLabel (Pion-Tonachini, Kreutz-Delgado, & Makeig, 2019). ICLabel provides the probability of each IC to be generated by brain, muscle, eye, heart, line noise, channel noise, and others. ICs were removed if the highest probability was generation by eye (if probability exceeded 25 %), muscle (if probability exceeded 90 %), and heart (if probability exceeded 60 %). These ICs were removed by setting their weight to 0. Across all tasks on average 5.71 ± 2.45 (median = 6) ICs were removed for each data set, ranging from 1 to 16. The resulting weight matrix was saved.

#### Pre-processing for time-frequency analysis

In a second step raw EEG data was pre-processed for analyzing time-frequency dynamics. Raw EEG data, segmented for the corresponding task, was high-pass filtered 0.1 Hz using a zero-phase Hamming windowed sinc FIR filter implemented in EEGLAB (Widmann et al., 2015). The same noisy channels, detected by the algorithm described above were removed, line noise was removed as described above, and data was re-referenced to average reference. To adjust the number of channels to the ICA weight matrix, the online reference Cz was restored before re-referencing and removed after-wards. The ICA matrix was imported, and EEG activity was re-calculated, with artefactual ICs removed. Finally rejected channels were interpolated using a spherical spline interpolation. Epochs were extracted by segmenting −1750 to 2000 ms around cue onset for cue-related analyses and −2950 to 2500 ms around target onset for target-related analyses. The larger time window for target-related analyses was due to having a baseline epoch prior to cue-onset and including the full response period. Artefactual epochs were rejected automatically using the same methods and parameters as described above. Moreover, an intra-individual outlier analysis was conducted on reaction times. This analysis was based on all correct trials. Reaction times were classified as outliers if their log-transformed and *z*-standardized form fell outside a threshold of ±3*SD* from the mean reaction time of an individual (Berger & Kiefer, 2021, 2023). These trials were rejected. Only trials with correct responses were kept for the timefrequency analyses. Overall, across switching tasks and cue- and target-related analyses, 20 ± 5 % (range: 9 – 42 %) of epochs were rejected: 5 ± 4 % due to errors, 1 ± 1 % due to reaction time outliers, and 15 ±4 % due to EEG artifacts. Finally, to achieve a similar signal-to-noise ratio between the conditions (switch and repeat), the number of trials per condition was equalized individually for each subject by randomly selecting *n* trials from the condition with the higher number of trials (where *n* is the number of trials in the condition with the lower number of trials).

#### Time-frequency analysis

To obtain the time-frequency representation of EEG power we calculated complex Morlet wavelets, adapting code provided by Cohen (2014, chapter 13). We extracted 25 frequencies logarithmically spaced between 3 and 60 Hz. The number of wavelet cycles changed as a function of frequency and ranged between 3 and 6. To account for edge artefacts, 1000 ms at the beginning and end of each epoch were discarded after wavelet convolution. For cue-related analyses, this resulted in epochs from −750 to 1000 ms relative to cue onset, with activity from −750 to −250 ms serving as baseline. For target-related analyses, epochs ranged from −1950 to 1500 ms relative to target onset, with activity from −1950 to −1450 ms serving as the baseline period. The target-related baseline period preceded cue onset regardless of cue presentation duration and ISI. Baseline activity was calculated across both conditions. We then averaged the mean instantaneous power over trials independently for each condition. The resulting time-frequency power spectrum was normalized to the baseline using decibel conversion. Finally, we downsampled the timefrequency representation from the initial sampling rate of 1000 Hz to 50 Hz.

#### Statistical Analyses

All statistical analyses were conducted with R (v3.6.3) (R Core Team, 2020) or MATLAB (R2021a, The MathWorks, Inc.), except otherwise specified. For mass-univariate analyses, we only included participants achieving a mean accuracy above or equal to 70 percent in the cognitive flexibility tasks. This led to the exclusion of of data from three participants in the parity/magnitude task, from two participants in the global/local task, and three participants in the number/letter task. No participant was excluded from all tasks. Additionally, we performed inter-individual outlier analyses for all manifest variables included in the structural equation models. Therefore, variables were *z*-transformed, after ensuring their approximate normal distribution (Hair, Black, Babin, & Anderson, 2019). A value was set to missing, if its absolute *z*-standardized form was larger than three. After the inter-individual outlier analysis, we *z*-standardized the variables again for further SEM analyses. We estimated the reliability of all manifest indicator variables in the structural equation models. We used Cronbach’s *α* for WMC and intelligence scores and Spearman-Brown corrected odd-even split correlations for the electrophysiological data.

#### Mass-univariate analyses with cluster-based permutation testing

To examine how instantaneous power in response to the cue and target differed between conditions, we used mass-univariate analyses (Groppe et al., 2011). For each voxel (representing EEG power of one time point, frequency, and electrode), we performed a parametric repeated measures *t*-test with the factor condition (switch vs. repeat). Significance level was set to *p* = .005 (Benjamin et al., 2018). We corrected for multiple comparisons using cluster-based permutation tests (Groppe et al., 2011; Maris & Oostenveld, 2007). Clusters are defined as significant voxels that are connected in time, frequency, and space. For this purpose, neighboring electrodes were defined as all electrodes that were directly adjacent to the seed electrode. Only voxels showing effects in the same direction were included in a cluster. The size of each empirical cluster was then compared to a distribution of cluster sizes derived from permuted data. Therefore, mass-univariate analyses were conducted on permuted data. In permuted data, the condition labels of trials have been randomly shuffled. For each permutation, the size of the largest cluster detected is extracted and contributes to the null distribution. We included 1,000 permutations and determined cluster significance by the joint examination of *t*-mass (sum of all *t* values within a cluster) and cluster size (amount of significant voxels within a cluster). In practice, these two criteria consistently yielded the same outcomes in the current study. The permutation test reveals whether a cluster of a specific extent is likely to be present in data where the condition labels of trials have been randomly shuffled (permuted data). If the extent of a cluster found in the original data is highly unlikely (*p <* .05) in permuted data, the empirical cluster is interpreted as representing meaningful condition differences. All other significant voxels are interpreted as chance findings. Here, significance level was set to *p* = .05, in order to explore all potentially relevant clusters (Benjamin et al., 2018). Importantly, cluster-based permutation tests neither reveal the exact time period when significant differences are present nor the exact frequency range or location of the detected effect, as they are based on probabilistic test statistics (Sassenhagen & Draschkow, 2019).

#### Structural equation modeling p

In a first step, we evaluated whether differences in event-related spectral perturbations (ERSP) between task conditions within a cluster do load onto a common flexibility factor across tasks. Therefore ERSP values within the core region of significant clusters were averaged and served as manifest variables in the structural equation models. Core regions were defined because cluster are subject to uncertainties regarding the spatial, temporal, and spectral extent (Sassenhagen & Draschkow, 2019). The core regions of the clusters were defined by the effect sizes 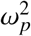 of the voxels contributing to the cluster. For this purpose, the effect sizes of a cluster were first logarithmized and then the mean plus one standard deviation was calculated as a cut-off value. Only those voxels whose effect size was greater than the cut-off value contributed to the core region. If this restriction caused clusters to split into different parts, only the largest sub-cluster was considered.

We modeled flexibility using a latent change model (McArdle, 2009; Schmitz & Krämer, 2023), as it addresses the issue of poor reliability of manifest difference scores. In line with previous research, the reliabilities of the cue-related manifest difference scores were found to be poor to unacceptable, with values ranging from .00 to .10 for theta and from .35 to .53 for alpha/beta in our data. Similarly, reliabilities of the target-related manifest difference scores reliabilities were found to be poor to unacceptable, with values ranging from .40 to .53 for alpha/beta. Because a latent change model captures the difference between two or more (repeated-measures) conditions on the latent level, it overcomes this critical limitation of manifest difference score. In the models we implemented, the two conditions of each task contribute to a switch and a repeat factor, respectively. Flexibility is modeled as the residual variance, when the switch factor is regressed on the repeat factor with a *β*-weight of 1. We added *m* − 1 method factors, where *m* − is the number of tasks. By including method factors, we account for task-specific associations between ERSPs. The use of *m* − 1 over *m* method factors is advantageous because it results in a globally identified model (Eid, 2000; Grayson & Marsh, 1994). Here, the parity/magnitude task served as the comparison standard, while we specified one method factor loading on all ERSP values measured in the global/local task and a second method factor loading on all ERSP values measured in the number/letter task. The method factors are thus defined as residual factors common to all ERSP values measured within the same task. Because all manifest variables had been *z*-transformed, the intercepts were set to zero.

In a second step, we combined the measurement models of flexibility into one single model. In addition to flexibility we modeled intelligence by using the four operation-related components of intelligence (processing capacity [PC], processing speed [PS], memory [M], and creativity [C]). Furthermore, WMC was modeled using the memory updating task, the operation span task, the sentence span task, and the location-letter binding task. With this model we investigated the latent correlations between cognitive flexibility, general cognitive abilities, and WMC.

All models were fitted with the R package *lavaan* (Rosseel, 2012), using the maximum likelihood algorithm. We used full information maximum likelihood (FIML) to account for missing data. We evaluated the goodness of fit of the models using the comparative fit index (CFI; Bentler, 1990) and the root mean square error of approximation (RMSEA; Browne & Cudeck, 1992). We considered a CFI of > .90 and a RMSEA of < .08 to indicate an acceptable fit of the model. A CFI > .95 and a RMSEA < .06 indicated a good model fit (Browne & Cudeck, 1992; Hu & Bentler, 1999). The two-sided critical ratio test was used to assess the statistical significance of the model parameters.

For the latent-change structural equation model (*df* = 162), with a sample size of *N* = 148, and an alpha error of *α* = .05, we can test the hypothesis of close fit (H0: *ε* ≤.05, H1: *ε*. ≥ 08; MacCallum, Browne, & Sugawara, 1996) with a power of 1 − *β* = .96. However, for the measurement models estimated in the first step (*df* = 12), the hypothesis of close fit can only be tested with low statistical power (1 − *β* = .25).

## Results

### Time-frequency dynamics in response to the cue

In all tasks and across conditions, the brain exhibited a general increase in theta power within the initial 400 ms following cue onset, most pronounced in the lateral occipital regions, and a general decrease in alpha/beta power beginning at 100 ms after cue onset (see first row of Figures 2a and 2c for frontal and occipital electrodes, respectively). The alpha/beta power decrease persisted throughout the remainder of the analysis period. It was descriptively strongest in the lateral occipital cortices. However, it was also present in the broader frontal region.

**Figure 2.**
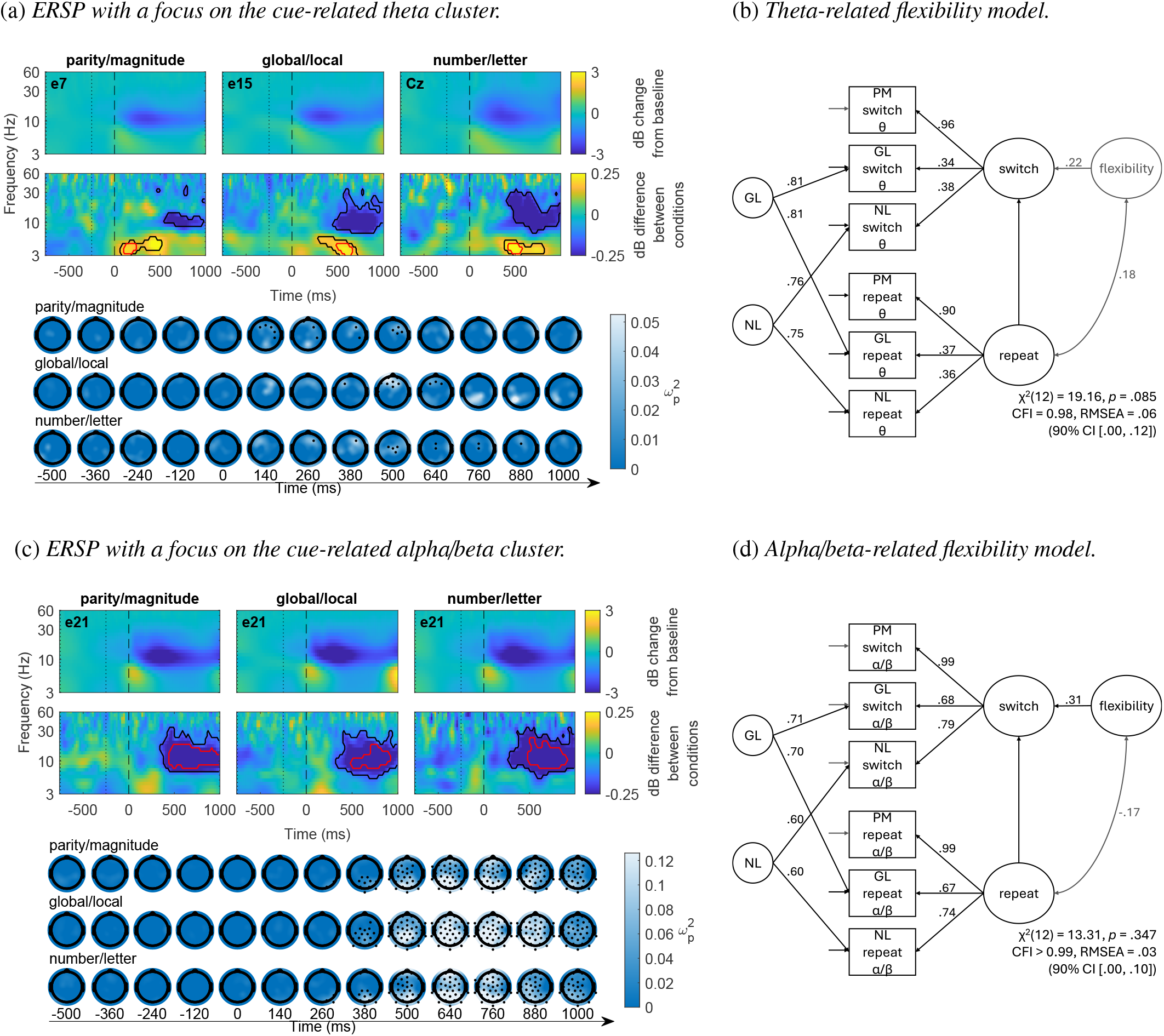
Event-related spectral perturbations in response to the cue (left columns) and latent-change models of cognitive flexibility (right columns). *Note*. **(a)** and **(c)** The first rows show the dB change of power in relation to the baseline (−750 to −250 ms relative to cue onset) averaged across both conditions. The name of the depicted electrode is shown in the top-left corner. The second rows display the dB difference between the conditions (switch minus repeat) at the same electrodes. Positive values represent greater power in switch compared to repeat trials, while negative values represent lesser power. The black border surrounds voxels belonging to a cluster, and the red border indicates voxels belonging to the core region of that cluster. The lower part of (a) and (c) shows the topographic distribution of effect sizes 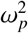, derived from univariate tests contrasting the two conditions, at an exemplary frequencies of (a) 5 Hz for the theta cluster and (c) 10 Hz for the alpha/beta cluster. Additionally, electrodes contributing to the respective clusters are depicted. **(b)** and **(d)** Latent-change models of cognitive flexibility based on mean power of the core region of the (b) theta and (d) alpha/beta cluster. The standardized path weights and correlation coefficients are shown next to the paths. Non-significant estimators are grayed out.

We found two distinct clusters across all three tasks in the empirical data, representing differences between conditions in theta power increase and alpha/beta power decrease in response to the cue (see second row of Figures 2a and 2c). Exact cluster statistics for each task can be found in Table A1 in the appendix.

There was a significantly greater theta increase in switch trials compared to repeat trials. Over the three tasks, this effect emerged at fronto-central electrodes roughly 300-550 ms after cue onset and explained about 3 % of the variance in theta power (lower part of Figure 2a). Condition differences were most pronounced 100 to 200 ms after cue onset for the parity/magnitude task and 500 to 600 ms after cue onset for the other two tasks, explaining up to 7 % of the variance (Table 1).

**Table 1.** Statistical description of the core regions of significant, cross-task generalizable clusters for cue- and target-related analyses.

| Task | $\omega_p^2$-cutoff | Frequency range | Time range | $t$-mass | cluster size | $\omega_p^2$ | | Contributing electrodes |
| --- | --- | --- | --- | --- | --- | --- | --- | --- |
| | | Hz | ms | sum | #voxel | mean ± SD | 95% CI | |
| cue-related theta cluster |  |  |  |  |  |  |  |  |
| parity/magnitude | >.036 | 3.4–4.4 | 110–230 | 98.91 | 27 | .042 ± .004 | [.040, .044] | 6 7 |
| global/local | >.045 | 3.0–4.9 | 470–650 | 229.48 | 56 | .052 ± .005 | [.050, .054] | 6 7 15 25 |
| number/letter | >.043 | 3.4–4.4 | 430–570 | 82.03 | 21 | .051 ± .006 | [.048, .054] | 3 Cz |
| cue-related alpha/beta cluster |  |  |  |  |  |  |  |  |
| parity/magnitude | >.088 | 7.2–28.4 | 390–990 | -6590.43 | 1047 | .121 ± .028 | [.119, .123] | 3 4 5 10 11 12 13 14 15 20<br>21 22 25 28 29 30 Cz |
| global/local | >.108 | 6.3–22.1 | 390–930 | -11250.80 | 1640 | .138 ± .027 | [.136, .140] | all electrodes except 8 18<br>26 27 |
| number/letter | >.091 | 8.1–25.0 | 310–910 | -6252.66 | 1028 | .119 ± .025 | [.117, .121] | all electrodes except 9 17<br>18 23 26 27 Cz |
| target-related alpha/beta cluster |  |  |  |  |  |  |  |  |
| parity/magnitude | >.067 | 8.1–28.4 | -310–490 | -7020.33 | 1293 | .092 ± .023 | [.090, .094] | 1 2 3 4 5 6 11 12 13 14 15<br>16 20 21 22 25 28 29 30<br>31 Cz |
| global/local | >.086 | 7.2–28.4 | -270–310 | -15431.97 | 2432 | .121 ± .029 | [.119, .123] | all electrodes except 8 18<br>26 27 |
| number/letter | >.072 | 7.2–25.0 | -330–610 | -8914.80 | 1641 | .097 ± .022 | [.095, .099] | 1 2 3 4 5 6 7 8 10 11 12<br>13 14 15 16 20 21 22 25<br>28 29 30 31 Cz |

*Note.* A core region is defined as voxels with an effect size greater than the mean plus one standard deviation of the corresponding cluster's effect sizes ( $\omega_p^2$ -cutoff).

In addition, there was a significantly greater decrease in alpha/beta power in switch trials compared to repeat trials. This difference was present throughout the cortex, but was most pronounced at left lateral parietal sites (lower part of Figure 2c). The difference was present approximately 100-200 ms after cue onset and persisted until the end of the extracted time period (1000 ms after cue onset). It explained about 6 % of the variance in alpha/beta power. The differences in condition were most evident between 400 to 900 ms after cue onset, explaining up to 25 % of the variance (Table 1).

### Testing for a common flexibility factor in cue-related power modulations

To test whether these power modulations load onto a common latent flexibility factor, we averaged the ERSP separately for each cluster’s core region, task, and condition. In a structural equation model, we used these averaged power values as indicators for the latent factor flexibility. See Table 1 for an overview of the core regions.

Table A2 in the appendix presents the reliabilities of the manifest indicator variables used in the structural equation models and their zero-order correlations. The ≥ reliabilities of the condition-specific ERSP values were found to be excellent for alpha/beta (≥.95) and mostly acceptable or good for theta (.65–.86).

In a first step, we tested whether the detected power changes load onto a common flexibility factor across tasks separately for each ERSP cluster. Specifically, we modeled flexibility as the latent difference between both task conditions using a latent change model to account for the unreliability of manifest difference scores. The latent change model with cue-related theta ERSP explained the data well, *χ*^2^(12) = 19.16, *p* = .085, CFI = .98, RMSEA = .06, 90% CI [.00, .12] (Figure 2b). However, flexibility could not be captured by this model, as the variance of the latent change factor was not significant (*p* = .517). Regarding the cue-related alpha/beta cluster, the latent change model provided a good explanation of the data, *χ*^2^(12) = 13.31, *p* = .347, CFI > .99, RMSEA = .03, 90% CI [.00, .10] (Figure 2d). Here, flexibility could be captured, as the variance of the latent change factor was significant (*p* < .001). Baseline alpha/beta power (i.e., in the repeat condition) was not associated with the magnitude of change (*r* = −.17, *p* = .175, 95% CI [−.40, .06]).

### Time-frequency dynamics in response to the target

Similarly to the cue-related power changes, in all tasks and across conditions, the brain exhibited a general increase in theta power around 400 to 500 ms following target onset, most pronounced at left and right lateral occipital electrode sites, and subsequently a general decrease in alpha/beta power lasting until 1000 ms post target for the parity/magnitude and number/letter task (first row of Figure 3a). For the global/local task, the alpha/beta decrease endured until 1500 ms post target. Not only was the alpha/beta decrease more persistent in the global/local task compared to both other tasks, also was the theta increase descriptively strongest in this task and visible throughout the cortex. The alpha/beta power decrease was descriptively strongest at lateral occipital electrode sites. However, it was also present in the broader frontal region.

**Figure 3.**
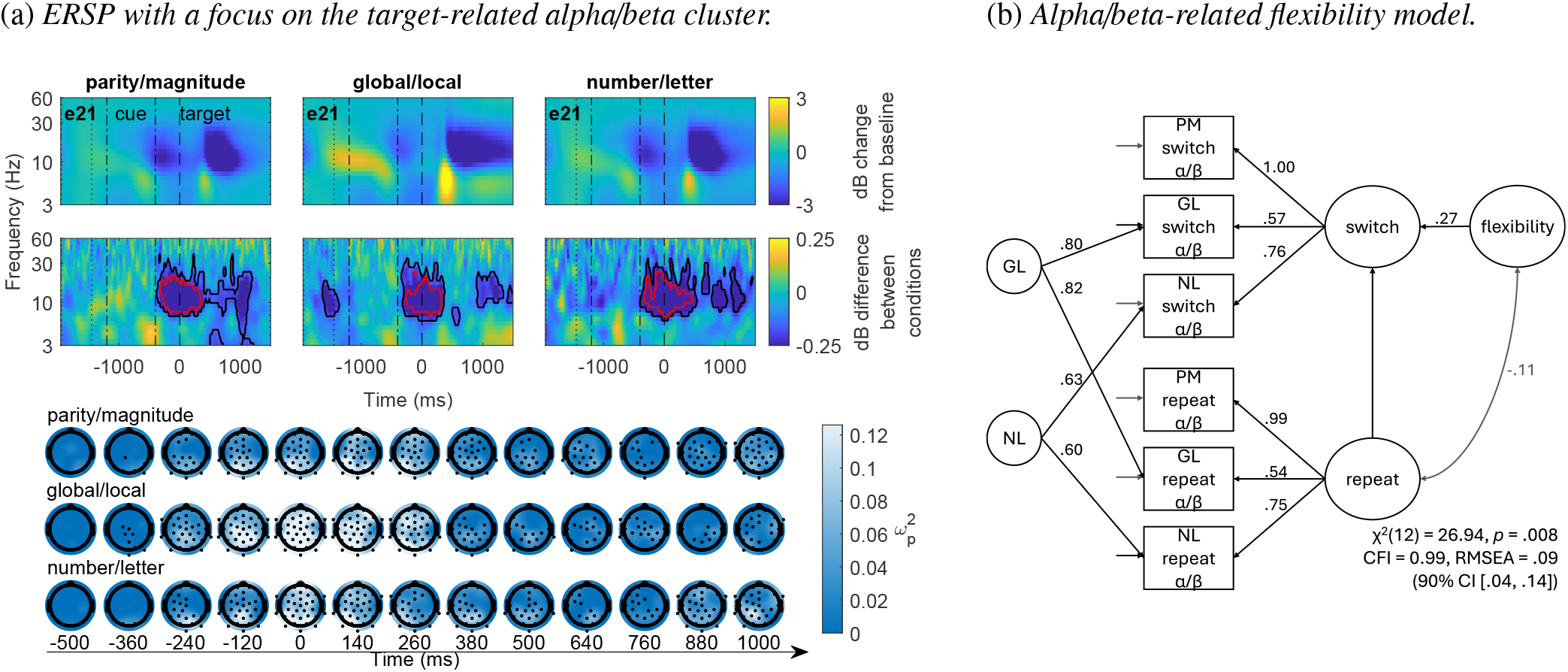
Event-related spectral perturbations in response to the target (left) and latent-change models of cognitive flexibility (right). *Note*. **(a)** The first row shows the dB change of power in relation to the baseline (−1950 to −1450 ms relative to target onset) averaged across both conditions. The dashed-dotted lines mark the period during which the cue was presented. The name of the depicted electrode is shown in the top-left corner. The second row displays the dB difference between the conditions (switch minus repeat) at the same electrodes. Positive values represent greater power in switch compared to repeat trials, while negative values represent lesser power. The black border surrounds voxels belonging to a cluster, and the red border indicates voxels belonging to the core region of that cluster. The lower part of (a) shows the topographic distribution of effect sizes 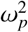 derived from univariate tests contrasting the two conditions, at an exemplary frequencies of 10 Hz for the alpha/beta cluster. Additionally, electrodes contributing to the respective cluster are depicted. **(b)** Latent-change model of cognitive flexibility based on mean power of the core region of the alpha/beta cluster. The standardized path weights and correlation coefficients are shown next to the paths. Non-significant estimators are grayed out.

Cluster-based permutation tests revealed three distinct clusters in the empirical data, representing differences between the switch and repeat conditions. As in response to the cue, conditions differed regarding the alpha/beta power decrease. In addition, statistical analysis of the data revealed a condition difference in a targetrelated theta increase and beta decrease. However, only the alpha/beta cluster generalized across all three tasks, while the theta and beta cluster did only emerge in the parity/magnitude and number/letter task, but not in the global/local task. Exact cluster statistics for all three clusters can be found in Table A1. The core region of the alpha/beta cluster is described in Table 1.

A significantly greater decrease in alpha/beta power occurred in switch trials compared to repeat trials in response to the target (second row of Figure 3a). This difference was present throughout the cortex, but was most pronounced at left lateral parietal electrode sites (lower part of Figure 3a). The difference emerged approximately 300 ms before target onset and persisted until the end of the extracted time period (1500 ms after cue onset). It explained about 6 % of the variance in alpha/beta power. The difference was most pronounced −300 to 500 ms around target onset and explained up to 26 % of the variance during this time window (Table 1). Alpha/beta effects are likely to be modulated by the tasks context prior to target onset, as participants prepare for the upcoming target after cue presentation and engage in preparatory processes up until the target can be processed.

The theta cluster manifested at right fronto-central sites, clearly before target onset for the number/letter task (−350 to 150 ms; core region: −270 to −70 ms) and with target onset for the parity/magnitude task (−210 to 490 ms: core region: 10 to 290 ms; Figure A2a in the appendix). The clear temporal onset before the target in the number/letter task suggests a methodological artifact, specifically a cue-related effect visible in this analysis epoch. Consequently, only for the parity/magnitude task, there is a significant target-related theta effect. Based on this pattern of findings, the target-related theta effect was not further considered as a general neural marker for cognitive flexibility.

A substantially larger decrease in beta power was observed in switch trials relative to repeat trials, indicating a significant relationship between target-related power changes and trial type. This difference was evident at right lateral central electrode sites between 700 and 1500 ms following target onset (Figure A2b). It accounted for approximately 4 to 5 % of the variance in beta power. The difference was most pronounced 1050 to 1400 ms after target onset, explaining up to 12 % of the variance during this time window. However, like the theta cluster, the beta cluster was only present in the parity/magnitude and number/letter task. Furthermore, the fact that the mean latency of condition differences in the beta cluster exceeded participants’ mean reaction time suggests that this cluster is unlikely to reflect target-induced cognitive flexibility processes. Consequently, this cluster, like the theta cluster, was not further tested as potential neural signature of cognitive flexibility.

### Testing for a common flexibility factor in target-related power modulations

In order to determine whether these power modulations load on a common latent flexibility factor, target-related power changes within the core region of the alpha/beta cluster were averaged. The mean was calculated independently for each task and condition. Comparable with the cue-related values, reliabilities of the averaged target-related condition-specific ERSP values (Table A2) were found to be excellent for alpha/beta (≥ .96).

The target-related latent change model converged and fit well to the target-related alpha/beta ERSP data, *χ*^2^(12) = 26.94, *p* = .008, CFI = .99, RMSEA = .09, 90% CI [.04, .14] (Figure 3b). Flexibility was captured by this model, as the variance of the latent change factor was significant (*p* < .001). Again, baseline alpha/beta power (i.e., in the repeat condition) was not associated with the magnitude of change: participants with higher baseline alpha/beta did not show significantly larger or smaller decreases than those with lower baseline alpha/beta (*r* = −.11, *p* = .390, 95% CI [−.35, .14]).

### Association between cognitive flexibility, intelligence, and working memory capacity

After determining suitable measurement models for flexibility, in a second step, we integrated the measurement models across clusters and explored their relation to general cognitive abilities and WMC. As evident from Table A2, WMC could be measured with good reliability (>.80), while for the BIS, with reliabilities ranging from .45–.75, only processing capacity had a sufficient reliability. Note that this is typical for the short form of the BIS. This is why it is recommended to only interpret the general ability factor, as we do (see Figure 4), rather than the operation-related manifest scores (Jäger et al., 1997).

**Figure 4.**
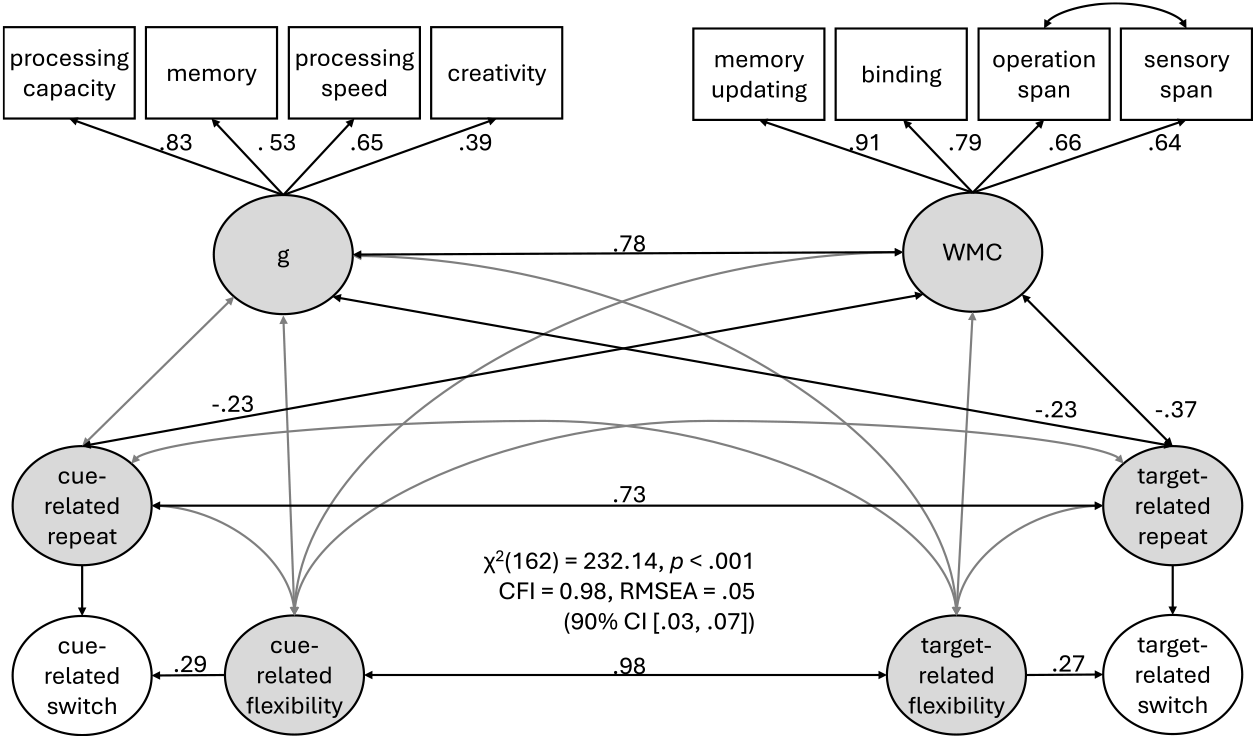
Combined latent-change models of cognitive flexibility based on alpha/beta power including intelligence (g) and WMC. *Note*. The standardized path weights and correlation coefficients are shown next to the paths. Non-significant estimators are grayed out.

We only included the two alpha/beta clusters in the integrated model, as these were capable of capturing common variance across the three flexibility tasks. In the integrated model, we allowed correlations between residual variances within condition-specific cue- and target-related activity (e.g., alpha/beta ERSP in the switch condition of the global/local task in response to the cue and the target). Furthermore, we allowed the residual variances of the sentence span and operation span tasks of the WMC battery to correlate, as these tasks are highly similar.

The model explained the data well, *χ*^2^(162) = 232.14, *p* < .001, CFI = .98, RMSEA = .05, 90% CI [.03, .07] (Figure 4). Cue-related and target-related flexibility could be captured by this model, as both latent change factors were significant (*p* < .001). The two latent flexibility factors were highly correlated (*r* = .98, *p* < .001, 95% CI [.95, 1.00]). The same was true for cue- and target-related alpha/beta power in the repeat condition (*r* = .73, *p* < .001, 95% CI [.65, .82]). Furthermore, cue- and target-related repeat factors, reflecting basic stimulus processing, showed significant latent correlations with WMC (cue: *r* = −.23, *p* = .016, 95% CI [−.40, −.05]; target: *r* = −.37, *p* < .001, 95% CI [−.53, −.20]) indicating a weak to moderate relationship. Additionally, the target-related, but not the cue-related, repeat factor exhibited a weak negative association with general cognitive abilities (*r* = −.23, *p* = .020, 95% CI [−.41, −.05]). Except for the substantial latent correlation between intelligence and WMC (*r* = .78, *p* < .001, 95% CI [.66, .89]), none of the other correlations were significant (−.17 ≤ *r* ≤ .16).

## Discussion

Our main goal was to identify oscillatory mechanisms underlying cognitive flexibility and assess their generalizability across tasks. We found that cognitive flexibility engages distinct neural oscillatory patterns: frontal theta increases reflect task-specific control processes, while parietal alpha/beta suppression represents domain-general attentional mechanisms. Critically, only alpha/beta effects generalized across tasks, supporting their role as neural markers of trait-like flexibility. These findings advance our theoretical understanding of flexibility by demonstrating that it involves sustained attentional processes spanning from cue presentation through target processing.

We first discuss the general oscillatory response to stimuli in cued switching tasks. Because these general response cannot disentangle basal from flexibility-specific processes, we then examine the condition-specific effects reflecting pure cognitive flexibility processes, and discuss their generalizability across tasks. Finally, we evaluate whether oscillatory responses can distinguish between cue- and target-related cognitive flexibility processes.

### General oscillatory response to task demands

Across all three cued task-switching tasks we find well known spectral perturbations in response to the cue and target: a transient theta increase followed by a sustained alpha/beta suppression. This pattern is present in both the lower-demand repeat condition and the higher-demand switch condition. The observed spectral perturbations were most pronounced in the occipital cortex, suggesting a visual cortical origin, consistent with the processing of visual stimuli.

Theta, alpha, and beta oscillations are not tied to single cognitive functions, but rather support a range of processes depending on their spectral, spatial, and temporal characteristics (Buzsáki, 2006; Vezoli et al., 2021). In case of visual processing, it has been shown that theta, alpha/beta, and gamma frequency bands function as channels, which route information between visual areas (Bastos et al., 2015; Michalareas et al., 2016). These studies find the same time-frequency patterns as we have observed here and have furthermore shown that theta and gamma frequencies are associated with feedforward and alpha/beta frequencies are associated with feedback influences. Activity within these frequencies may reflect the reallocation of attention by feedforward and feedback loops, possibly ensuring that the task can be completed successfully (Sadaghiani & Kleinschmidt, 2016).

### Condition-specific effects: flexibility-related neural modulations

To isolate flexibility-specific processes, we contrasted switch versus repeat trials. This revealed two distinct oscillatory signatures in the theta and alpha/beta frequency ranges. These signatures differed from the general oscillatory response in terms of their spatial and temporal characteristics, and their generalizability across tasks.

### Theta as task-specific cognitive control

This contrast revealed a significantly stronger theta power increase in frontal regions during switch trials in response to the cue. The frontal theta enhancement can be linked to a large body of evidence associating mid-frontal theta activity with cognitive control processes, such as conflict monitoring, task-set updating, and rule switching (Cavanagh & Frank, 2014; Cavanagh & Shackman, 2015). Our findings thus replicate previous findings (e.g. Capizzi et al., 2020; Cooper et al., 2017) and support the notion that cognitive control is exerted to a greater extent in switch trials than in repeat trials. However, we should emphasize that we did not observe strictly mid-frontal theta effects. In our switching tasks, we observed effects that were more right-frontal than mid-frontal. Notably, the temporal and spatial maximums of these effects differed between tasks. Nevertheless, the stronger cue-related theta increase in switch trials compared to repeat trials was present in all three shifting tasks. While mid-frontal theta activity is typically associated with cognitive control processes and is thought to originate primarily from midcingulate cortex (Cavanagh & Shackman, 2015), the right lateral prefrontal cortex has also been strongly implicated in cognitive control, particularly in response inhibition (Aron, Robbins, & Poldrack, 2004). According to Ridderinkhof, Ullsperger, Crone, and Nieuwenhuis (2004), mid-frontal control signals are translated into action plans in the lateral prefrontal regions. Supporting this view, Verbruggen, Aron, Stevens, and Chambers (2010) experimentally showed that the right ventral inferior frontal cortex is associated with updating action plans. Further evidence for the involvement of lateral prefrontal regions in executive functions comes from a meta-analysis on working memory tasks, requiring some sort of executive control. This analysis revealed that mental manipulations, such as updating or reordering information, are consistently associated with increased activation in right lateral prefrontal cortex (Wager & Smith, 2003). However, there is also metaanalytic evidence questioning the specific involvement of the lateral frontal cortex in switching, linking this region primarily with executive working memory (Wager, Jonides, & Reading, 2004).

Regarding the target, we found a greater theta increase in switch trials only for the parity/magnitude task. The task-specific nature of theta effects likely reflects the distinct cognitive demands across switching tasks. The number/letter and global/local tasks allow preparatory filtering of relevant information (Ravizza & Carter, 2008). In contrast, the parity/magnitude task requires reactive control to resolve conflicts arising from the automatic processing of both stimulus categories. This explains why only the parity/magnitude task showed significant target-related theta effects: Interference must be resolved when the target appears, necessitating continued cognitive control.

### Alpha/beta as domain-general attention

Switch trials also showed stronger alpha/beta suppression in comparison to repeat trials, maximal in the left parietal association cortex. This suppression occurred after the initial occipital response and persisted from cue through target processing. Alpha/beta suppression indicates attentional resource allocation (Van Diepen et al., 2019; Sauseng et al., 2006).

The left parietal focus of this condition-specific alpha/beta effect underscores its role in attentional control and selective attention (Peylo, Hilla, & Sauseng, 2021; Behrmann, Geng, & Shomstein, 2004). Notably, this top-down attentional modulation was not limited to the initial cue processing, as around target onset, stronger left parietal alpha/beta suppression was observed in switch trials compared to repeat trials, mirroring the effect seen for the cue. Given that condition differences emerged several hundred milliseconds before target onset and persisted for an equivalent duration after onset, it is reasonable to assume that subjects maintain attention for a prolonged time after the cue is presented. This suggests that they still require more attentional resources to process the target stimulus when the rule-set changes between trials.

These dynamics are consistent with the notion that reallocation of attention, elevated working memory demands, and the goal-directed regulation of cognition and behavior engage distributed control systems, including medial frontal and lateral parietal areas (Sadaghiani & Kleinschmidt, 2016; Wager et al., 2004). Together, these theta and alpha/beta effects support the engagement of fronto-parietal control networks during task switching (Chang et al., 2023), with theta reflecting transient control signals and alpha/beta reflecting sustained attentional adjustments.

### Cross-task generalizability of neural flexibility markers

A main goal of our analyses was to establish whether individual differences in the observed condition-specific effects reflect cognitive flexibility beyond task-specific patterns. We addressed this using structural equation modeling to extract latent factors representing shared variance across switching tasks.

Frontal theta effects, despite being present in all tasks, showed no significant shared variance across tasks. This indicates that while cognitive control is engaged universally, the specific control mechanisms are tailored to each task’s demands. Each switching task requires different forms of attention shifting—between stimulus dimensions (parity/magnitude), between perceptual levels (global/local), or between stimuli (number/letter).

In contrast, alpha/beta suppression exhibited significant common variance across all tasks for both cue- and target-related activity. This provides the first direct evidence for cognitive flexibility as a latent construct that generalizes beyond task-specific demands. The emergence of this common factor validates the convergent validity of these tasks as measures of cognitive flexibility.

In a second step, we combined the two models of cue- and target-related flexibility, both of which were based on oscillatory alpha/beta band activity, into a single model. In addition, we added two other broad performance constructs, namely, intelligence and WMC, in order to relate cognitive flexibility to these. The analysis revealed that cue- and target-related cognitive flexibility reflect a highly overlapping process, sharing 96 % of their variance, suggesting they index the same underlying construct. In addition, alpha/beta suppression in the baseline (repeat) condition was positively correlated across cue- and target-related measures, sharing 53 % of the variance.

However, neither flexibility factor correlated significantly with intelligence or WMC, supporting previous findings of dissociation between flexibility and general cognitive ability (Kopp, Maldonado, Scheffels, Hendel, & Lange, 2019; Santarnecchi et al., 2021). Nevertheless, empirical findings on the relationship between flexibility, intelligence, and WMC are inconsistent and may furthermore depend on the specific neuronal or behavioral indicators used (Schmitz & Krämer, 2023).

Baseline alpha/beta suppression (repeat condition) showed weak negative correlations with intelligence and WMC, consistent with the broader literature on alpha suppression and cognitive abilities (Hilger, Spinath, Troche, & Schubert, 2022). Importantly, flexibility measures were independent of baseline suppression, indicating that individual differences in flexibility are distinct from general processing efficiency.

Taken together, we find that alpha/beta suppression is attributable to a general flexibility factor, which is distinct from other performance measures, such as intelligence and WMC. A central contribution of the present study lies in the extraction of common variance across three cued switching tasks, enabling us to identify whether a shared latent process underlies the observed spectral perturbation patterns. While a considerable portion of variance in the neural data is accounted for by task-specific effects, our results provide, to our knowledge, the first direct evidence that cognitive flexibility can be modeled as a latent process that generalizes across distinct tasks.

### Disentangling cue- and target-related processes

The high correlation between cue- and target-related flexibility factors raises questions about whether they represent distinct processes. Several lines of evidence suggest they reflect a single, sustained attentional process rather than separate mechanisms.

Both effects occurred in the same frequency range and location. Critically, cue-related suppression persisted until analysis epoch end, while target-related effects began several hundred milliseconds before target onset. This temporal overlap indicates that attention reallocation initiated by the cue continues through target processing, rather than representing separate preparatory and execution phases.

This pattern supports two-stage theories of task switching (Kiesel et al., 2010), which propose that cues trigger goal-shifting and rule retrieval, while targets require stimulus-response mapping. Our findings suggest that attentional processes initiated during goal-shifting must be maintained through target processing, explaining why residual switch costs persist even with preparation time.

From a neural network perspective, this sustained attention likely reflects the need to maintain activation of task-relevant networks while suppressing competing rule sets (Miller & Cohen, 2001). Repeat trials benefit from pre-activated neural patterns representing the current rule set, while switch trials require active reconfiguration of network states, which is a process that cannot be completed before target appearance (Kiesel et al., 2010; Qiao, Zhang, Chen, & Egner, 2017).

Conversely, switch and repeat trials do not differ in terms of the need to activate the correct stimulus-response mapping and retrieve the appropriate motor plan. However, the ease with which this happens should depend on the pre-activation of rule sets and stimulus-response mappings, and therefore on trial type (Jost, Baene, Koch, & Brass, 2015; Qiao et al., 2017). In line with this, we find no evidence that the categorization of the target activates different processes in case of repeating or switching.

In summary, our findings suggest that upon presentation of the cue, more cognitive resources are needed when participants have to switch the current rule-set in comparison when repeating it. This shows up due to the stronger alpha/beta suppression in switch trials. The increased activation continued until the presentation of the target and beyond. This indicates that attention allocation and the adaption of attentional weights of relevant networks was started by the cue and endured until the target occurred (Meiran, 2000).

### Limitations and future directions

While the present study advances our understanding of the neural underpinnings of cognitive flexibility, it also has some limitations.

Our aim of examining the spectral, spatial, and temporal specificity of the oscillatory effects underlying cognitive flexibility was partially compromised by our focus on the 3–60 Hz frequency range, excluding higher gamma frequencies (>60 Hz). Gamma activity, in particular, reflects localized neural network activations (Fries, 2015; Buzsáki, 2006) and may benefit from high-density EEG recordings. With only 32 electrodes, our study lacks the spatial resolution necessary to precisely localize effects, especially for task-specific phenomena like the cue-related theta ERSP. Future research should employ high-density recordings optimized for gamma analysis and a high spatial resolution. Despite these limitations, our data-driven approach confirmed previous findings and emphasizes the importance of theta and alpha/beta, underscoring the robustness of these bands in cognitive flexibility.

Cluster-based permutation testing, while controlling false positives, may reduce sensitivity to smaller effects (Noble, Scheinost, & Constable, 2020). Complementary hypothesis-driven analyses and advanced methods like threshold-free cluster enhancement (Smith & Nichols, 2009) could increase sensitivity while maintaining statistical rigor.

Lastly, although we employed three distinct tasks to assess cognitive flexibility and are able to generalize from task-specific demands to the abstract construct of flexibility, future work should investigate whether these findings are specific for flexibility. Previous research has shown that an oscillatory response in the form of a transient theta increase and sustained alpha/beta suppression is not unique to flexibility tasks, but rather is involved in general conscious, goal-directed processing (e.g. Xiang, Huang, Luo, Ma, & Zhang, 2021).

### Conclusion and outlook

This study advances our understanding of cognitive flexibility’s neural foundations through several key contributions. We identified and confirmed oscillatory patterns linked to cognitive flexibility. We identified alpha/beta suppression as a robust neural marker of domain-general flexibility, while demonstrating that theta effects reflect task-specific control processes. The methodological framework employed, combining datadriven time-frequency analyses with structural equation modeling, not only increases confidence in the robustness of previous findings but also refines them. Building on the assumption that neuronal communication relies on frequency-specific channels, which are influenced by structural and spatial network properties, we found support for the notion that lower frequencies are associated with top-down attention control, highlighting their importance for flexible, goal-directed behavior. The sustained nature of flexibility-related alpha/beta suppression provides novel evidence for twostage models of task switching, showing that attentional processes must remain active from cue through target processing.

These insights have profound implications for understanding individual differences in cognitive flexibility and its independence from general intelligence. By establishing alpha/beta suppression as a neural signature of domain-general flexibility, this work provides a neurobiological foundation for cognitive flexibility as a distinct construct. This framework opens new avenues for assessing flexibility capacity, developing targeted interventions, and understanding why some individuals excel at adapting to changing demands while others struggle. Most importantly, our findings demonstrate that cognitive flexibility—a capacity fundamental to learning, problem-solving, and adaptive behavior—can be measured and understood at the neural level. This represents a crucial step toward bridging the gap between cognitive theory and neurobiology, establishing that the brain’s oscillatory dynamics provide a window into one of our most essential cognitive abilities.

## Acknowledgements

This research was funded by the Deutsche Forschungsgemeinschaft (DFG, German Research Foundation) – 442137702; 399695734. The funders had no role in study design, data collection and analysis, decision to publish or preparation of the manuscript.

## Author Contributions

M. J. H.: Formal analysis, Methodology, Validation, Visualization, Writing – original draft, Writing – review & editing; C. L.: Conceptualization, Data curation, Investigation, Project administration, Writing – review & editing; A.-L. S.: Conceptualization, Funding acquisition, Methodology, Project administration, Supervision, Writing – original draft, Writing – review & editing;

## Appendix

### Supplementary tables and figures

**Figure A1.**
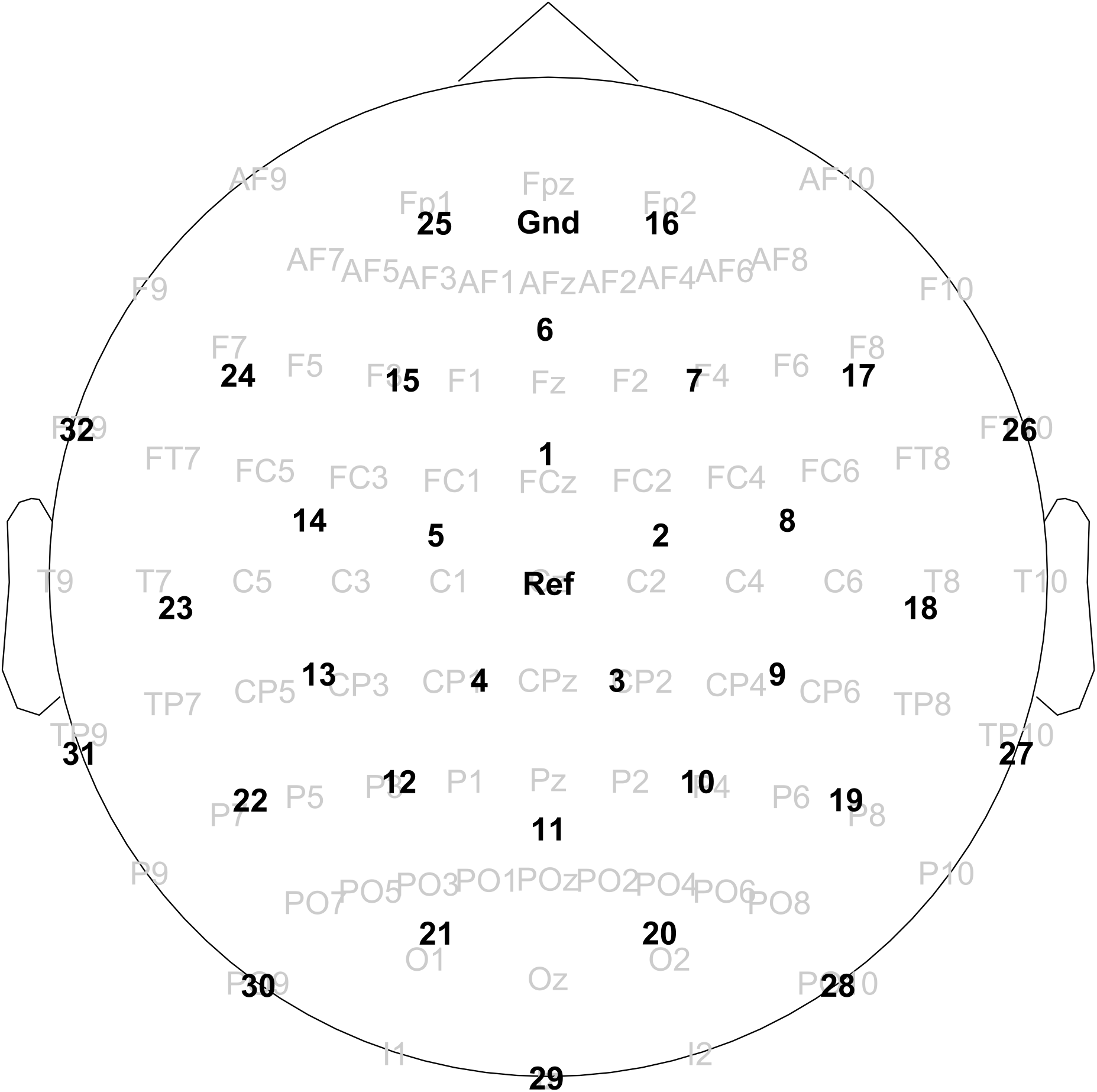
Equidistant electrode layout used in the current study. *Note*. The 10-10 % electrode layout is depicted in gray in the background. Cz served as online reference. The figure was created using the Fieldtrip (Oostenveld, Fries, Maris, & Schoffelen, 2011) layout files.

**Table A1.**
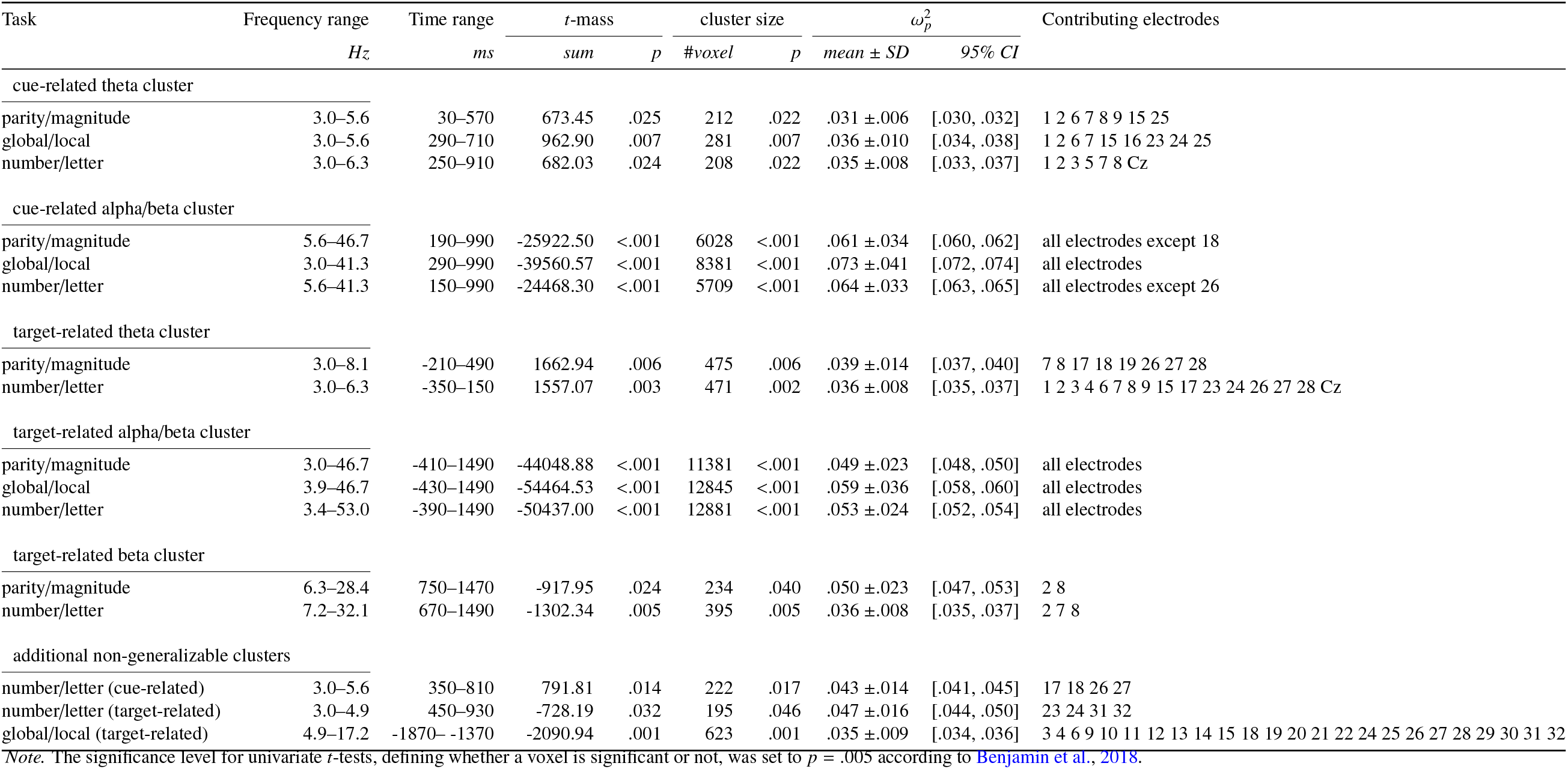
Statistical description of significant clusters for cue- and target-related analyses.

**Table A2.** This table shows the correlations between the manifest variables separately for both conditions (switch and repeat) with their corresponding reliabilities on the diagonal.

| Measure | 1 | 2 | 3 | 4 | 5 | 6 | 7 | 8 | 9 | 10 | 11 | 12 | 13 | 14 | 15 | 16 | 17 | 18 | 19 | 20 | 21 | 22 | 23 | 24 | 25 | 26 |
| --- | --- | --- | --- | --- | --- | --- | --- | --- | --- | --- | --- | --- | --- | --- | --- | --- | --- | --- | --- | --- | --- | --- | --- | --- | --- | --- |
| 1 WMC - memory updating | .88 |  |  |  |  |  |  |  |  |  |  |  |  |  |  |  |  |  |  |  |  |  |  |  |  |  |
| 2 WMC - binding | .69 | .82 |  |  |  |  |  |  |  |  |  |  |  |  |  |  |  |  |  |  |  |  |  |  |  |  |
| 3 WMC - operation span | .58 | .47 | .89 |  |  |  |  |  |  |  |  |  |  |  |  |  |  |  |  |  |  |  |  |  |  |  |
| 4 WMC - sentence span | .61 | .32 | .66 | .87 |  |  |  |  |  |  |  |  |  |  |  |  |  |  |  |  |  |  |  |  |  |  |
| 5 BIS - processing capacity | .60 | .54 | .38 | .41 | .75 |  |  |  |  |  |  |  |  |  |  |  |  |  |  |  |  |  |  |  |  |  |
| 6 BIS - memory | .34 | .33 | .23 | .27 | .43 | .58 |  |  |  |  |  |  |  |  |  |  |  |  |  |  |  |  |  |  |  |  |
| 7 BIS - processing speed | .42 | .44 | .20 | .22 | .50 | .43 | .49 |  |  |  |  |  |  |  |  |  |  |  |  |  |  |  |  |  |  |  |
| 8 BIS - creativity | .13 | .22 | .09 | .13 | .35 | .13 | .34 | .45 |  |  |  |  |  |  |  |  |  |  |  |  |  |  |  |  |  |  |
| 9 cue-related theta (PM) - switch | .05 | .09 | .12 | .11 | .15 | .11 | .08 | .11 | .86 |  |  |  |  |  |  |  |  |  |  |  |  |  |  |  |  |  |
| 10 cue-related theta (PM) - repeat | .02 | .02 | .12 | .14 | .14 | .11 | .05 | .17 | .84 | .81 |  |  |  |  |  |  |  |  |  |  |  |  |  |  |  |  |
| 11 cue-related theta (GL) - switch | .18 | .22 | .11 | .14 | .16 | .10 | .21 | -.08 | .31 | .29 | .79 |  |  |  |  |  |  |  |  |  |  |  |  |  |  |  |
| 12 cue-related theta (GL) - repeat | .20 | .21 | .14 | .11 | .11 | .09 | .12 | -.13 | .33 | .33 | .78 | .82 |  |  |  |  |  |  |  |  |  |  |  |  |  |  |
| 13 cue-related theta (NL) - switch | .09 | .20 | .00 | .09 | .13 | .06 | .12 | -.03 | .35 | .29 | .39 | .42 | .78 |  |  |  |  |  |  |  |  |  |  |  |  |  |
| 14 cue-related theta (NL) - repeat | .07 | .23 | .01 | .07 | .13 | .02 | .08 | .01 | .32 | .29 | .35 | .35 | .70 | .65 |  |  |  |  |  |  |  |  |  |  |  |  |
| 15 cue-related alpha/beta (PM) - switch | -.19 | -.14 | -.11 | -.12 | -.11 | -.16 | -.12 | -.20 | .12 | .03 | .17 | .02 | .14 | .08 | .97 |  |  |  |  |  |  |  |  |  |  |  |
| 16 cue-related alpha/beta (PM) - repeat | -.18 | -.14 | -.15 | -.15 | -.11 | -.15 | -.09 | -.14 | .09 | .03 | .19 | .05 | .13 | .05 | .93 | .96 |  |  |  |  |  |  |  |  |  |  |
| 17 cue-related alpha/beta (GL) - switch | -.14 | -.16 | -.02 | -.07 | -.04 | -.22 | -.19 | -.18 | .09 | .05 | .15 | .03 | .08 | -.04 | .70 | .67 | .96 |  |  |  |  |  |  |  |  |  |
| 18 cue-related alpha/beta (GL) - repeat | -.17 | -.19 | -.05 | -.11 | -.04 | -.22 | -.12 | -.19 | .05 | .02 | .15 | .06 | .01 | -.15 | .64 | .68 | .94 | .96 |  |  |  |  |  |  |  |  |
| 19 cue-related alpha/beta (NL) - switch | -.16 | -.13 | -.02 | -.05 | -.08 | -.04 | -.11 | -.17 | .27 | .17 | .19 | .05 | .21 | .13 | .77 | .72 | .60 | .53 | .96 |  |  |  |  |  |  |  |
| 20 cue-related alpha/beta (NL) - repeat | -.12 | -.14 | -.05 | -.01 | -.02 | .00 | -.05 | -.11 | .23 | .16 | .20 | .07 | .23 | .11 | .68 | .72 | .59 | .57 | .91 | .95 |  |  |  |  |  |  |
| 21 target-related alpha/beta (PM) - switch | -.30 | -.25 | -.08 | -.12 | -.15 | -.17 | -.15 | -.25 | .12 | .04 | .11 | .00 | .08 | .05 | .76 | .66 | .59 | .56 | .54 | .46 | .97 |  |  |  |  |  |
| 22 target-related alpha/beta (PM) - repeat | -.31 | -.29 | -.13 | -.16 | -.17 | -.17 | -.12 | -.19 | .10 | .04 | .13 | .01 | .06 | .03 | .69 | .71 | .57 | .59 | .50 | .50 | .95 | .96 |  |  |  |  |
| 23 target-related alpha/beta (GL) - switch | -.15 | -.18 | -.07 | -.10 | -.10 | -.27 | -.19 | -.06 | -.01 | -.04 | -.01 | -.09 | -.04 | -.06 | .40 | .42 | .78 | .75 | .36 | .39 | .55 | .59 | .98 |  |  |  |
| 24 target-related alpha/beta (GL) - repeat | -.16 | -.21 | -.09 | -.11 | -.09 | -.27 | -.13 | -.07 | -.04 | -.05 | .01 | -.06 | -.09 | -.13 | .31 | .37 | .68 | .75 | .27 | .34 | .50 | .57 | .95 | .97 |  |  |
| 25 target-related alpha/beta (NL) - switch | -.20 | -.15 | .06 | -.11 | -.10 | -.10 | -.14 | -.24 | .15 | .11 | .04 | -.04 | .07 | .05 | .59 | .48 | .50 | .45 | .69 | .57 | .73 | .66 | .48 | .41 | .97 |  |
| 26 target-related alpha/beta (NL) - repeat | -.18 | -.17 | .01 | -.07 | -.09 | -.11 | -.12 | -.24 | .12 | .09 | .06 | -.01 | .15 | .05 | .55 | .51 | .54 | .52 | .64 | .67 | .71 | .70 | .55 | .51 | .91 | .96 |

*Note.* Correlations are calculated on $N = 139$ for memory updating and sentence span, $N = 133$ for binding, $N = 135$ for operation span, $N = 140$ for BIS, $N = 142$ for parity/magnitude task, $N = 144$ for global/local task, and $N = 134$ for number/letter task. Reliabilities were estimated with Cronbach's $\alpha$ for WMC and BIS and with Spearman-Brown corrected odd-even split correlations for the electrophysiological data. WMC: working memory capacity; BIS: Berlin intelligence structure test; PM: parity/magnitude task; GL: global/local task; NL: number/letter task;

**Figure A2.**
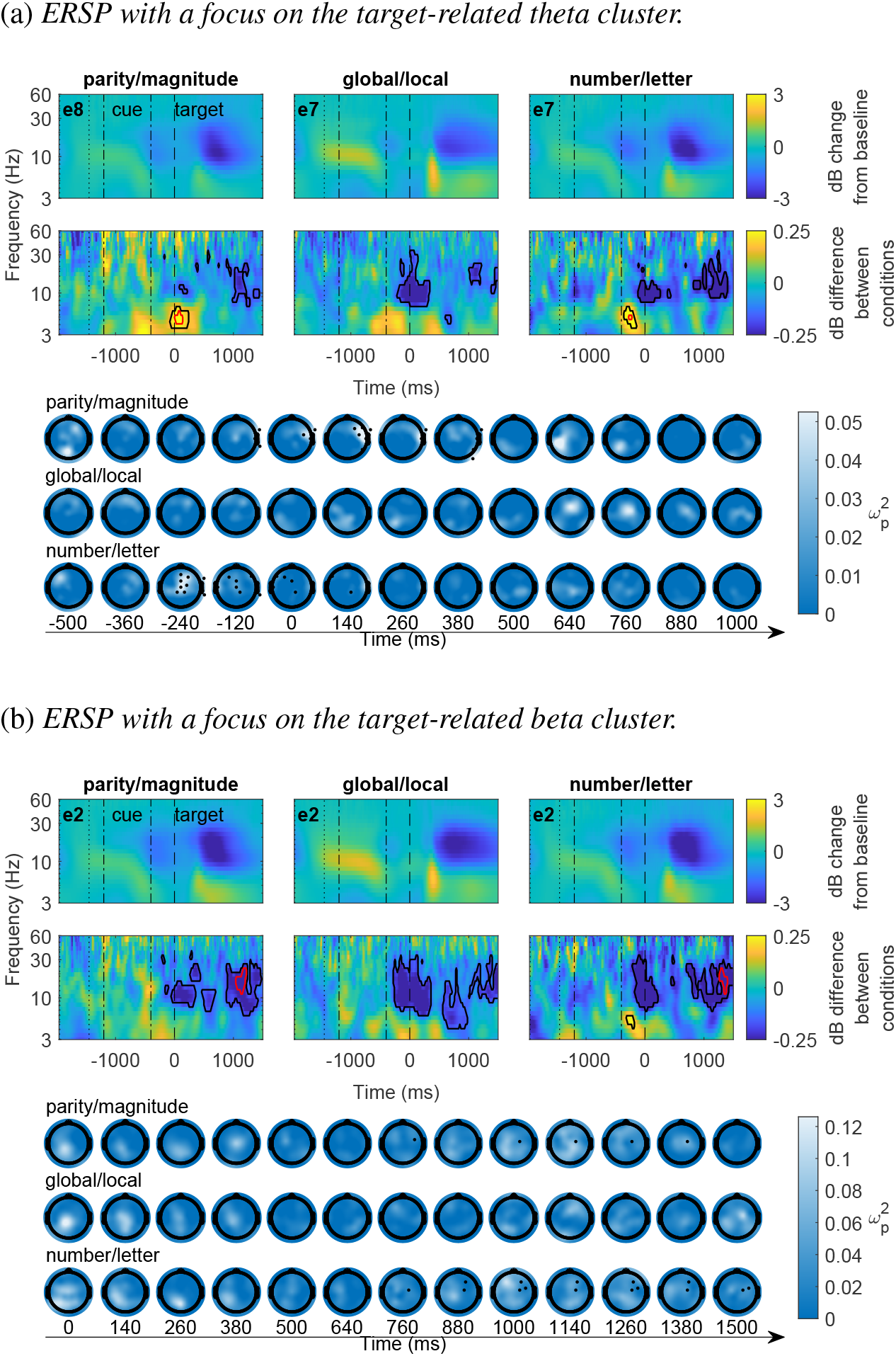
Event-related spectral perturbations in response to the target for non-generalizable clusters. *Note*. The first rows of (a) and (b) show the dB change of power in relation to the baseline (−1950 to −1450 ms relative to target onset) averaged across both conditions. The dashed-dotted lines mark the period during which the cue was presented. The name of the depicted electrode is shown in the top-left corner. The second rows display the dB difference between the conditions (switch minus repeat) at the same electrodes. Positive values represent greater power in switch compared to repeat trials, while negative values represent lesser power. The black border surrounds voxels belonging to a cluster, and the red border indicates voxels belonging to the core region of that cluster. The lower part of (a) and (b) shows the topographic distribution of effect sizes 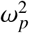, derived from univariate tests contrasting the two conditions, at an exemplary frequencies of (a) 5 Hz for the theta cluster and (b) 20 Hz for the beta cluster. Additionally, electrodes contributing to the respective clusters are depicted.

